# Intra- and extra-cellular environments contribute to the fate of HIV-1 infection

**DOI:** 10.1101/2020.07.22.195719

**Authors:** Sneha Ratnapriya, Miranda Harris, Angela Chov, Vladimir Vrbanac, Maud Deruaz, Joseph Sodroski, Alon Herschhorn

**Affiliations:** Division of Infectious Diseases and International Medicine, Department of Medicine, University of Minnesota, Minneapolis, Minnesota 55455; Humanized Immune System Mouse Program, Ragon Institute of MGH, MIT and Harvard and Center for Immunology and Inflammatory Disease; Department of Immunology Cancer and Virology, Dana-Farber Cancer Institute, Boston, MA 02215, USA; Department of Microbiology, Harvard Medical School, Boston, MA 02215, USA; Institute for Molecular Virology, University of Minnesota, Minneapolis, Minnesota 55455, USA

## Abstract

HIV-1 entry into host cells leads to one of three alternative fates: 1) HIV-1 elimination by restriction factors, 2) establishment of HIV-1 latency, or 3) active viral replication in target cells. Here we developed an improved system for monitoring HIV-1 fate and provide evidence for the differential contribution of the intracellular environment as well as extracellular environment found in organs of BLT humanized mouse to the fate of HIV-1 infection.

## MAIN

Cells containing a latent HIV-1 provirus are rare in people living with HIV (PLWH) and only a small fraction of these cells carry a replication-competent provirus capable of replenishing the viral population upon HIV-1 reactivation.^1,2^ Very low number of latent cells makes the identification and isolation of these cells extremely difficult in samples from PLWH.^3,4^ Thus, several model systems have been developed in recent years to investigate and provide new insights into the establishment and maintenance of the latent HIV-1 reservoir.^5,6^ A recent and popular system uses a reporter HIV-1-based vector in which one fluorescent protein is transcribed from the HIV-1 LTR to monitor HIV-1-mediated transcription and a second fluorescent protein is transcribed from an internal promoter to report HIV-1 entry and provirus integration in infected cells.^7,8^ Here, we used similar building blocks to construct a system, designated HI.fate, with improved elements to simultaneously monitor HIV-1 replication and latency at single-cell and population levels (Fig. 1a).^9^ Our reporter system is based on the NL4-3 backbone and maintains genetic stability as a result of differences between the upstream and downstream long terminal repeat (LTR) sequences of the viral genome.^10^ These differences significantly decrease potential recombination events between the two LTRs and the excision of the provirus, which has been previously reported to potentially limit the use of the system.^11^ Two fluorescent proteins were selected for optimal sensitivity in primary cells: E2-Crimson, which is transcribed from the HIV-1 LTR and serves as a marker of HIV-1 gene expression, and ZsGreen, which is transcribed from an internal promoter and serves as a marker for HIV-1 entry and integration into the host genome. The two fluorescent proteins are bright and exhibit favorable and well-separated excitation/emission profiles that allow measurement of the fluorescence intensity by separate lasers without the need for compensation.^12,13^

**Figure 1.**
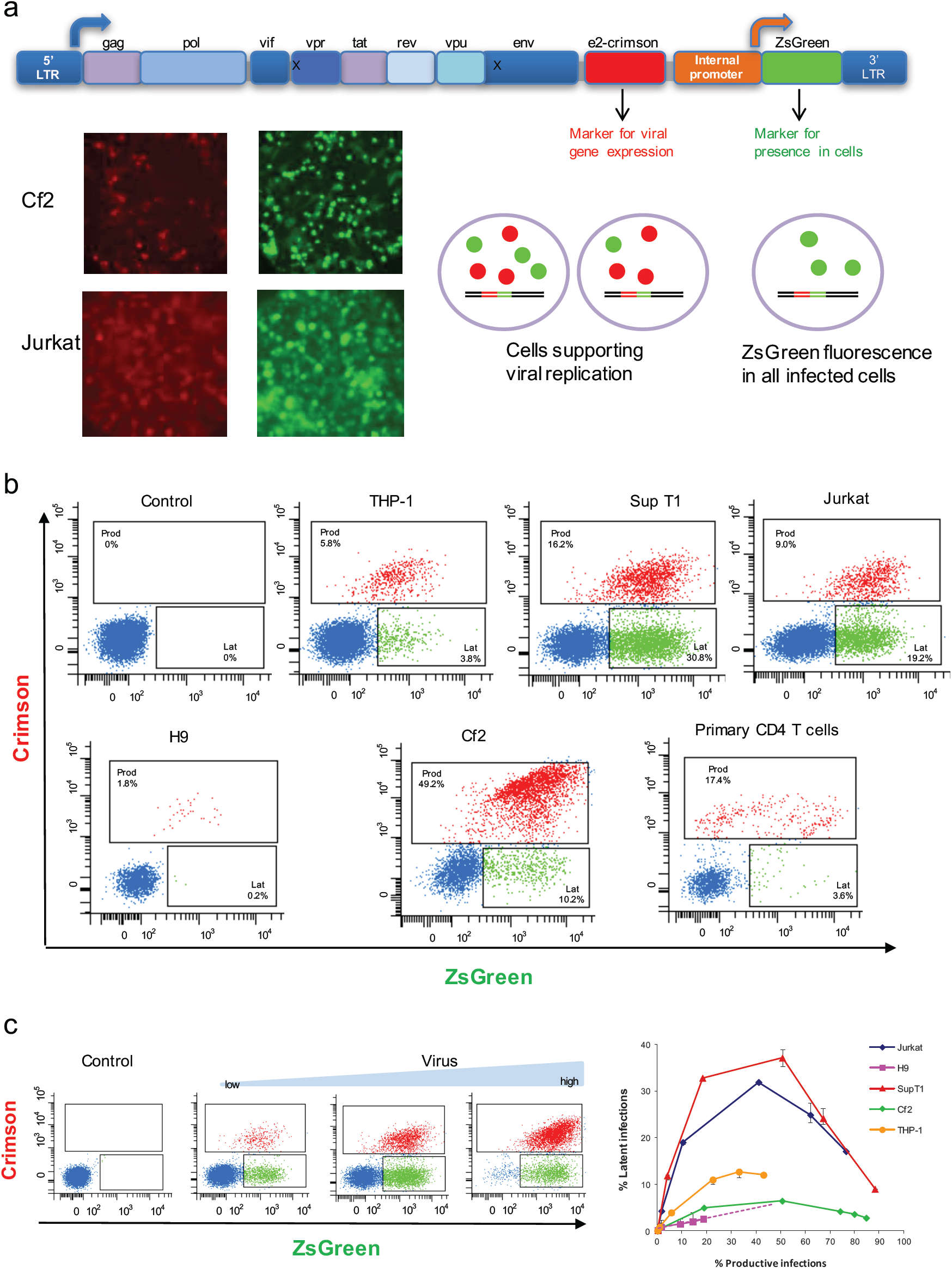
An improved HIV-1 reporter vector (HI.fate) distinguishes alternative fates of HIV-1 infection in target cells. (**a**) HI.fate is based on the chimeric molecular clone HIV-1_NL4-3_, which contains the 5’-long terminal repeat (LTR) from the NY5 isolate and the 3’-LTR from the LAV isolate. Differences between the two LTR sequences decrease the frequency of undesirable recombination events during vector manipulations. Lower panels – infection of different cells by HI.fate.E (EF1a-HTLV chimeric internal promoter) viruses leads to the expression of E2-Crimson and ZsGreen fluorescent proteins, which are visualized under a fluorescence microscope. (**b**) Flow cytometric analysis of different cells infected by HI.fate.E viruses. THP-1 cells were differentiated using 100nM PMA; primary CD4 T cells were isolated from peripheral blood mononuclear cells (PBMC) by negative selection and activated with anti CD3/CD28 beads prior to infection. Cells that support HIV-1 gene expression are shown in red and those supporting HIV-1 latency are shown in green. Unifected cells are colored blue (**c**) Increasing amounts of HI.fate.E viruses were used to infect target cells. The left panel shows the readout for 10,000 cells. Fluorescence profile of titered viruses was plotted for each cell type.

To analyze the fate of HIV-1 infection in different cells, we cloned the EF1a-HTLV chimeric promoter as an internal promoter into our reporter vector (HI.fate.E), prepared viruses pseudotyped with VSV-G envelope, and infected several T cells, including primary CD4+ T cells, as well as Cf2Th, canine thymic epithelial cells, and THP-1 (Fig. 1). Infected cells were analyzed by two-color flow cytometry 48 hours (cell lines) or 96 hours (primary CD4 T cells) post-infection to measure the frequency of latent cells versus cells that support HIV-1 gene expression. We observed significant variation in the fate of HIV-1 infection in the different target cells. To analyze this variability on a comparable scale, we titered the viruses on each type of cell and plotted the fluorescence profile of each cell type to assess the distribution of the latent and “replicating” phenotypes over an extended range of infection (we use HIV-1-mediated gene expression as a marker for replication even though HIV-1 is not fully replicating in these cells) (Fig. 1c). Notably, SupT1 and Jurkat cells supported high levels of HIV-1 latency whereas in other cells such as H9, Cf2Th and activated primary CD4+ T cells, active HIV-1 replication was reproducibly more dominant. Since the integration sites at the population level are not expected to be significantly distinct among the different cells, and we infected the different cells with the same reporter viruses, it is likely that the different phenotypes are related to the unique cellular environment of each cell type. At high multiplicity of infection, the frequency of cells that support HIV-1 replication was increased. This is expected because active HIV-1 LTR-mediated transcription has a dominant phenotype in our system (i.e. cells infected with two viruses - one latent and the second actively expressing the E2-Crimson - are phenotypically identified as supporting HIV-1 replication).

To optimize our reporter system and increase the sensitivity of detection of infected cells, we tested alternative internal promoters to mediate ZsGreen expression (Supplementary Fig. 1). We also fused the ZsGreen and GFP via the self-cleavage peptide 2A to express a self-cleaved fusion protein that leads to simultaneous expression of two green fluorescence proteins. HI.fate.SFFV showed minimal interference of viral transcription from the upstream LTR promoter (Supplementary Fig. 1 panels b and c) and it was selected for *in vivo* studies in a humanized BLT (bone marrow-liver-thymus) mouse model. We infected 4 humanized mice intravenously with the HI.fate.SFFV virus pseudotyped with the VSV-G envelope protein every other day for 7 days to ensure efficient infection (Fig. 2). The 4 humanized mice were generated from 2 different human donors (n=2 per donor). On day 7 we sacrificed the mice and collected different organs for flow cytometric analysis. The fate of HIV infection was evaluated by measuring the green and far red fluorescence in live, CD4+CD45+ cells present in different tissues. When we collectively analyzed cells from all organs of all four mice, we detected a significant (*P* value = 0.009) higher proportion of latent cells than cells supporting HIV-1 replication in our BLT mouse model (Fig. 2c). Within each of these groups (replicating vs latent), we observed statistically significant differences when comparing the specific phenotype in different organs (e.g. comparing latent cells in thymus and latent cells in blood; Fig. 2c). The spleen was a major source of cells supporting replication, with a comparable number of these cells and latent cells (*P* value > 0.99; Fig. 2d). Differences between the two populations were most notable in the blood, although for as yet unknown reasons, the number of positive cells isolated from the blood was significantly lower than the number of cells collected from other tissues. Specifically, for cells isolated from the blood, the number of latent cells was significantly higher than the number of cells supporting replication, but statistical significance was observed only for one-tail analysis (one-tail *P* value = 0.03). We observed a similar pattern in cells isolated from the lungs, where the difference between the two populations was smaller but the variability of the results among the mice was much lower. The number of latent cells in the lymph nodes and thymus was not statistically different than the number of cells supporting HIV-1 replication in these organs (Fig. 2d). Overall, these results point to possible variations in the fate of HIV-1 infection in different organs, characterized by a different extracellular environment. A complex relationship between the intra- and extra-cellular environments would determine the network of cellular molecules in the cells and contribute to their interactions with HIV-1 proteins and viral replication pathways. Our observations are consistent with and add to previous knowledge on the nature of HIV-1 replication. Previous quantification of the number of HIV-1 proviruses in the reservoir of PLWH documented higher numbers of resting CD4+ T cells containing HIV-1 provirus in PBMC than in the lymph nodes.^1^ As resting CD4+ T cells support no or low HIV-1 replication, an increased number of proviruses is expected to result in a dominant latency state in infected cells in the blood compared to those in the lymph nodes, as observed in our study. Moreover, high HIV-1 latency in the blood and extensive replication in the lymph nodes are well-documented during the clinically asymptomatic period of HIV-1 replication in PLWH and non-human primate models.^14,15^ We analyzed HIV-1 fate in four humanized BLT mice; larger experiments with increased number of animals may be needed to obtain higher statistical significance of the observed differences among different organs and to provide a higher resolution analysis. Nonetheless, our study provides important insights into the parameters that contribute to the fate of HIV-1 infection.

**Figure 2.**
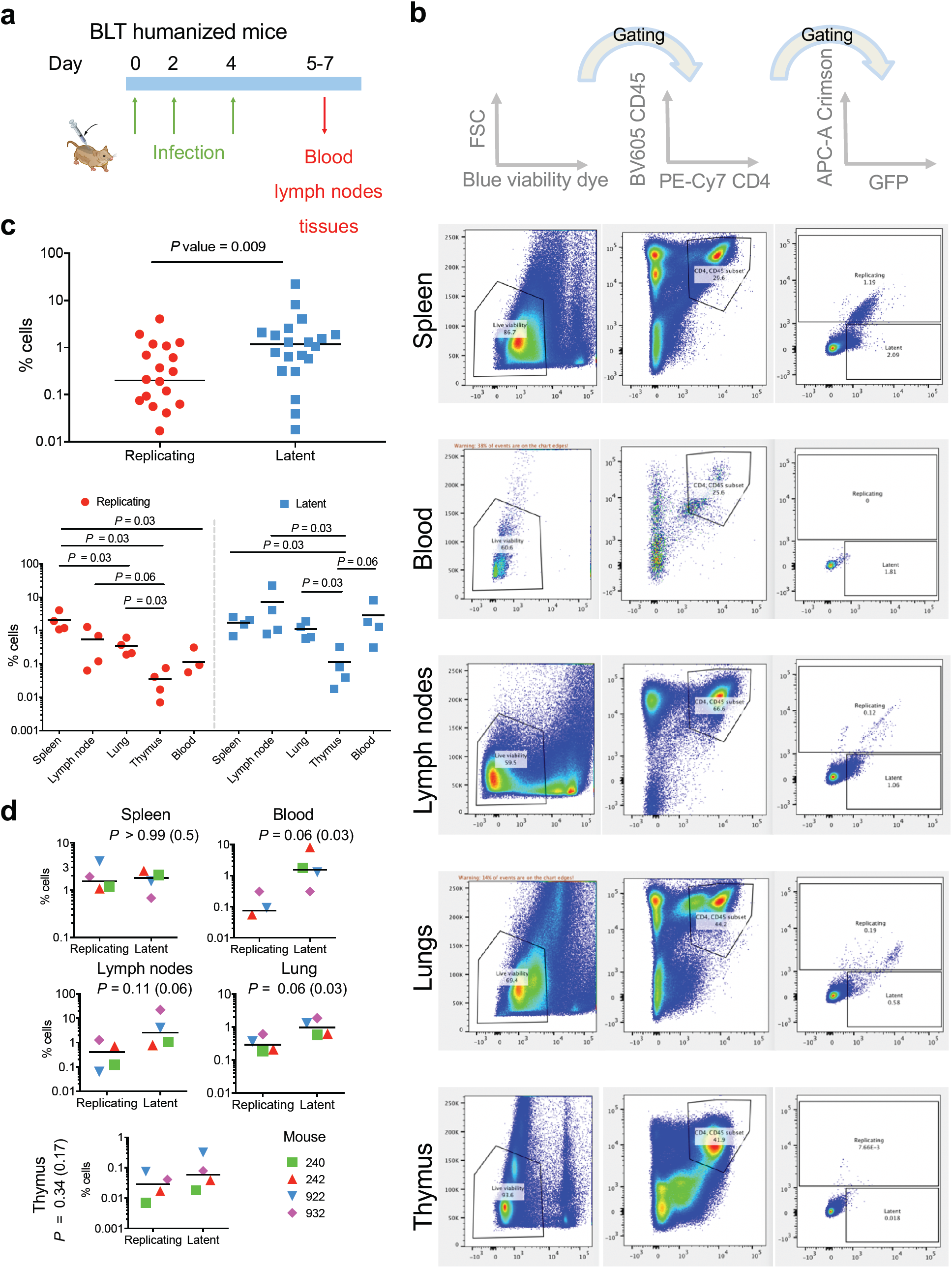
Fate of HIV-1 infection in BLT humanized model. **a.** Schematic layout of the experiment. **b.** Flow cytometric analysis of GFP and Crimson expression in live, CD4+ and CD45+ cells from different organs of a single BLT humanized mouse. Cell populations were identified and gated, and identical gates were applied to all tissues and mice. **c.** Statistical analysis of the differences between the overall number of latent cells and those supporting HIV-1 replication (top panel) and of the differences between the number of cells with specific phenotype in different organs (bottom panel) of four BLT humanized mice. **d.** Statistical analysis of the differences between the number of latent cells and those supporting HIV-1 replication for each organ. For panels **c** and **d**, *P* is two-tail *P* value of Mann-Whitney test and, when used, a value in parenthesis is a one-tail *P* value.

## METHODS

### Plasmid Construction

All HI.fate vectors are based on the HIV-1 pNL4-3.HSA.R-E- plasmid backbone, which was obtained from Nathaniel Landau through the NIH AIDS Reagent Program, Division of AIDS, NIAID, NIH (Cat# 3417). To construct HI.fate.E, three DNA fragments containing the 1) *e2-crimson* gene, 2) EF1a-HTLVI promoter, and 3) *zs-green* gene, were amplified by PCR. Sequences required for Gibson Assembly were added to the fragments during amplification and the three DNA fragments were cloned into the pNL4-3.HSA.R-E-plasmid, which was digested with Not I and Xho I restriction enzymes (NEB), using Gibson Assembly (NEbuilder; NEB). All subsequent HI.fate vectors containing different internal promoters were constructed by digesting HI.fate.E with BsiW I and Sac II restriction enzymes (NEB) and cloning each promoter, which was synthesized as a gene block (IDT), into the digested vector using Gibson Assembly (NEbuilder; NEB). HI.fate vectors expressing the Zs-Green protein fused to EGFP through the 2A self-cleavage peptide were constructed by digesting HI.fate.E with BsiW I and Xho I restriction enzymes (NEB) and cloning two fragments: 1) zs-green gene, and 2) 2A-egfp gene, which were amplified by PCR, into the digested vector using Gibson Assembly (NEbuilder; NEB).

### Production of Recombinant HI.fate viruses

We produced viruses as we previously described^16–19^ with the exception that we transfected 293T cells with two plasmids instead of the three plasmids routinely used for pseudovirus preparation. HI.fate (different versions) and a plasmid expressing VSV-G or HIV_KB9_ envelope glycoproteins were cotransfected at the mass ratio of 9:1 (9 Hi.fate / 1 Env) using Effectene (Qiagen) according to the manufacturer’s instructions. After a 48-hour incubation, the cell supernatant was collected and centrifuged for 5 minutes at 600-900x g at 4°C. The amount of p24 in the supernatant was measured using the HIV-1 p24 antigen capture assay (Cat# 5421, Advanced BioScience Laboratories) and the virus-containing supernatant was frozen in single-use aliquots at −80°C.

### Viral Infection Assay

A single-round infection assay was performed in 24-well or 6-well plates by spinoculation or by adding the specified amount of p24 (or increasing amounts for virus titration) to the cells. The cells were incubated for 48 or 72 hours and analyzed by flow cytometry.

### Animal Care

Humanized mouse experiments were performed according to NIH guidelines for the housing and care of laboratory animals. Protocols were reviewed and approved by the Institutional Animal Care and Use Committee (IACUC) of Massachusetts General Hospital and the University of Minnesota.

### Generation of humanized BLT mice and in vivo infection

Female BLT-NOD-scid IL2Rg^−/–^ (NSG) mice (The Jackson Laboratory) were housed in a pathogen-free facility at Massachusetts General Hospital and reconstituted with human tissue as described.^20^ Briefly, sublethally irradiated mice were transplanted under the kidney capsule with 1 mm^3^ fragments of human fetal liver and thymus, and injected intravenously with purified CD34^+^ human fetal liver cells to generate humanized BLT mice. Human immune cell reconstitution was monitored after surgery by measuring the concentration of human CD4+ T cells in the peripheral blood by flow cytometry. BLT mice showed no clinical signs of GvHD at any time during the experiment. Mice were infected 3 times intravenously at day 0, 2 and 4 with HI.fate viruses in a total volume of 500 μl PBS. Animals were euthanized 5 to 7 days after the first HIV infection and tissues were harvested for flow cytometry analysis.

### Flow cytometry

Human lymphocyte characterization was performed on an LSR II (BD Biosciences). Lungs from BLT mice were minced using scissors and incubated in medium containing Liberase (0.02 mg/mL) and DNAse (5 mg/ml) (both from Sigma-Aldrich). Spleen, liver, thymus and lymph nodes were minced using scissors. Processed tissues were passed through a 40 μm mesh strainer to obtain single-cell suspensions. Remaining red blood cells were removed using RBC lysis buffer (Sigma-Aldrich). Cells were stained using the anti-human monoclonal antibodies fluorophore-conjugated (Biolegend) Brilliant Violet 605™ anti-human CD45 (HI30), and PE/Cy7 anti-human CD4 Antibody (clone RPA-T4). Dead cells were excluded using LIVE/DEAD ™ Fixable Blue Dead Cell Stain Kit (Life Technologies). Cells were fixed with 2% paraformaldehyde after staining. Whole blood (100 μl) was directly stained with antibodies and treated with FACS lysing solution (BD Biosciences) before analysis. CountBright™ Absolute Counting Beads (ThermoFisher) were used to determine cell numbers.

## MATERIALS AVALIABILITY

Our system was distributed to several research groups in USA, Canada, and Europe and is available upon request.

## AKNOWLEDGMENTS

We thank NIH AIDS Reagent Program for providing the pNL4-3.HSA.R-E-plasmid. A.H. is the recipient of a phase II amfAR research grant (109285-58-RKVA) for independent investigators. This work was supported by the Harvard University Center for AIDS Research, the Humanized Immune System Mouse Program at Ragon Institute of MGH, MIT and Harvard, and internal funds of the Department of Medicine at the University of Minnesota.

## Notes

### Competing Interest Statement

The authors have declared no competing interest.

