## Supplementary Figure 1 for "Intra- and extra-cellular environments contribute to the fate of HIV-1 infection"

### Supplementary Information

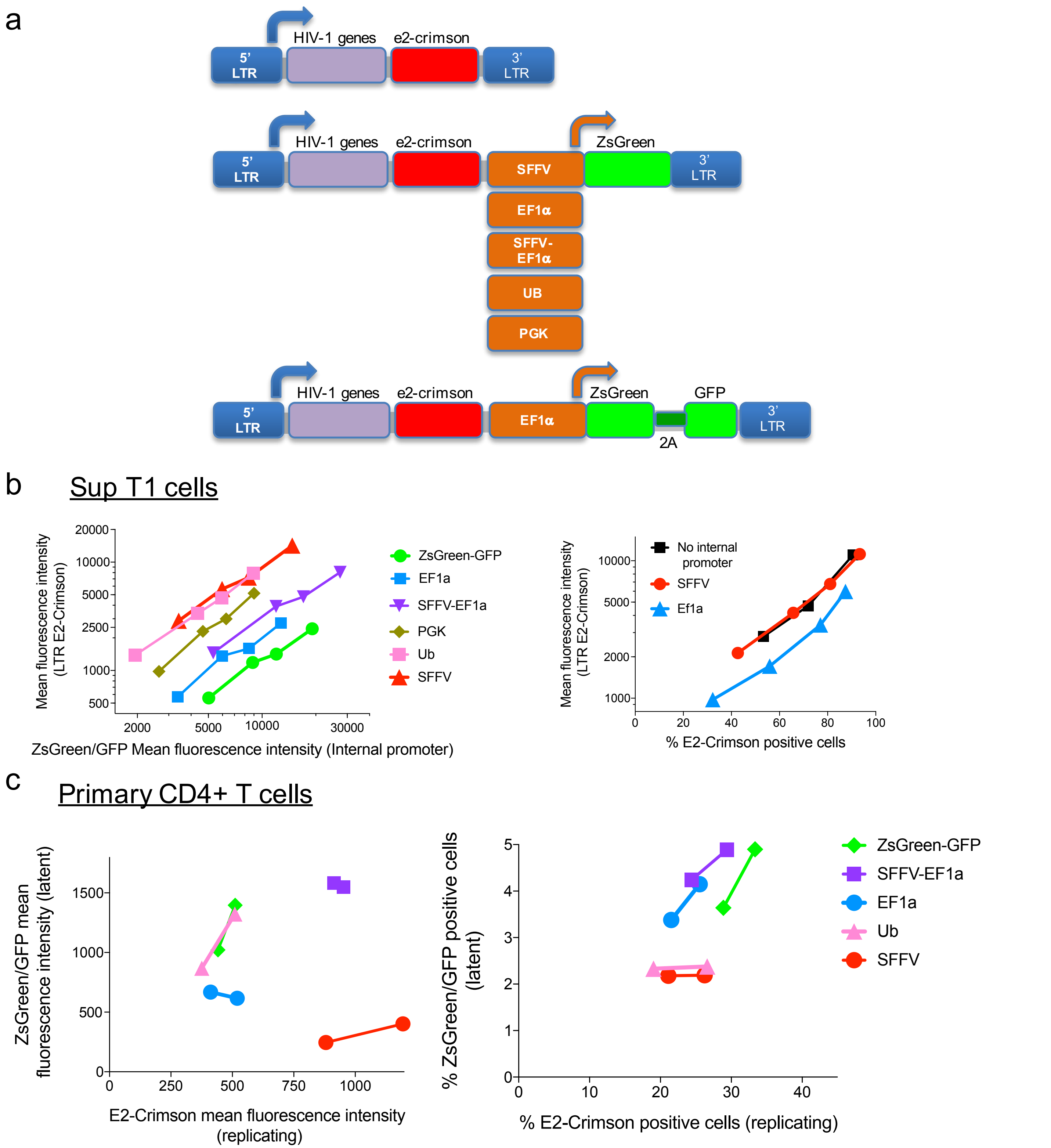


**Supplementary Figure 1. The effects of internal promoters and fused green fluorescent proteins on HIV-1 transcription.** (**a**) A schematic representation of the different vectors that were tested in the study. HI.fate was engineered to include the specified internal promoters or the ZsGreen fused to GFP via the 2A self-cleavage peptide for simultaneous expression of two green fluorescence proteins and increased sensitivity of detection. (**b**) Sup T1 cells were infected with increasing amounts of HI.fate viruses that were prepared by using the different vectors shown in **a**. Left – mean fluorescence intensity (MFI) of LTR-mediated expression versus MFI of internal promoter-mediated expression for different internal promoters. Right – MFI and frequency of cells supporting HIV-1 gene expression. Note that HI.fate.SFFV exhibit similar E2-Crimson as a vector with no internal promoter, indicating that the SFFV promoter does not significantly interfere with the upstream LTR-mediated transcription. (**c**) Primary CD4 T cells were isolated from PBMC by negative selection and activated with anti CD3/CD28 beads prior to infection. Cells were infected with 20ng or 100ng p24 of the indicated HI.fate viruses. Left – relationship between the MFI of latent cells and cells supporting HIV-1 gene expression for different internal promoters; Right – similar to the left panel but the % positive cells were plotted.
